# An Eco-Evolutionary Modelling Framework for Mosquito Host Specialisation

**DOI:** 10.64898/2026.08.25.747012

**Authors:** Alexander Sadykov, Dina Sadykova, Tina Mukherjee, Ben Matthews, Joao Marques, Mario Recker

**Affiliations:** The Centre for Ecology and Conservation, University of Exeter, UK; The Centre for Ecological and Evolutionary Synthesis, University of Oslo, Norway; UK Centre for Ecology and Hydrology, Wallingford, UK; Institute for Stem Cell Science and Regenerative Medicine, GKVK, Bangalore, India; Department of Zoology, University of British Columbia, Vancouver, Canada; Institut de Biologie Molèculaire et Cellulaire du CNRS, Strasbourg, France; Institute for Tropical Medicine, University of Tübingen, Tübingen, Germany

## Abstract

Host preference varies widely across mosquitoes, with many species feeding opportunistically on diverse vertebrate hosts, while others show strong fidelity to specific hosts. *Anthropophilia*, the behavioural preference for feeding on humans, is a defining characteristic of some mosquito species responsible for the transmission of major human diseases, including malaria, dengue, and yellow fever. The evolution of *anthropophilia* therefore has profound epidemiological implications because increased human biting elevates vectorial capacity and disease transmission potential. However, the ecological and evolutionary mechanisms driving this extreme specialisation has not yet been fully elucidated and remain difficult to unify across laboratory and field studies. Here we present an eco-evolutionary modelling framework that links genetically determined mosquito traits with spatially structured host environments. Our framework integrates innate olfactory sensitivity, blood meal-derived fitness benefits, and spatio-temporal host accessibility. Two complementary indices are introduced: a *local specialisation index*, capturing short-term ecological feeding strategies, and a *co-evolutionary index*, capturing long-term genetic coupling between host detection and resource utilisation. Our results demonstrate that host specialisation is not a default evolutionary outcome but an environmentally gated process, which is favoured in resource-poor or temporally varying habitats and strongly filtered by seasonality. The framework yields testable predictions regarding when specialisation emerges, persists, or collapses, with direct implications for predicting vector-borne disease risk in changing environments.

## 1 Introduction

### 1.1 Host Specialisation

Mosquitoes (family *Culicidae*) exhibit a range of host-feeding behaviours, from generalism, i.e. opportunistic feeding on multiple vertebrate classes, to specialism and strong preference for specific hosts, such as mammals or birds (reviewed e.g. in [1]). While most of the 3,500 known species of mosquito are generalists or specialise on non-human vertebrates, some mosquitoes, such as *Aedes aegypti* a.e.*Anopheles gambiae*, have evolved extreme preference for humans, referred to as anthropophilia. This has profound consequences for the epidemiology of vector-borne diseases, such as dengue, yellow fever or malaria, because their transmission is maximised when vectors concentrate bites on competent hosts and avoid non-infectious blood meals [2]. Consequently, understanding the drivers of mosquito host choice is central to predicting and managing vector-borne disease risk.

Anthropophilic behaviour has evolved repeatedly across diverse mosquito taxa and is believed to have emerged through ecological interactions between ancestral mosquitoes and expanding human populations, especially under climatic shifts. For example, recent genomic modelling suggests that the anthropophilic *Ae. aegypti* specialist diverged from its generalist forest ancestor approximately 5,000 years ago. This corresponds to the end of the African Humid Period, when the drying of the Sahel forced mosquitoes into human settlements, where stored water provided stable breeding sites and high densities of humans provided abundant resources for effective reproduction [3, 4, 5, 6] (McBride et al. 2014a; Rose et al. 2020; Metz et al. 2023; Rose et al. 2023). Work by (Rose et al. 2020) also showed that variation in preference for human odour across different *Ae. aegypti* populations in sub-Saharan Africa was strongly associated with dry season intensity and human density, supporting the hypothesis of an evolutionary transition towards anthropophilia rather than behavioural plasticity.

It is well established that host specialism is underpinned by specific genetic signatures predominantly related to odour sensing [3, 7] (McBride et al. 2014; McBride 2016). A key example is *Ae. aegypti*, whose domestic form *Ae. aegypti aegypti*, which has spread globally and almost exclusively bites humans, is behaviourally distinct from the ancestral forest form (*Ae. aegypti formosus*), which inhabits the tropical and subtropical regions of Africa and is primarily a zoophilic and opportunistic feeder. This behavioral shift has been shown to correlate with elevated expression of the *AaegOr4* odorant receptor gene, making them highly sensitive to volatiles in human skin and therefore underlies stronger attraction to humans vs. other animals[3] (McBride et al. 2014b). Equally, genomic analyses of different ecotypes of *Culex pipiens* have identified multiple regions containing chemosensory genes that are associated with preferences for avian vs. mammalian hosts (e.g.[8, 9, 10] (Fritz et al. 2015; Bell et al. 2024; Noreuil and Fritz 2021)). This genetic variability provides the foundation upon which selective forces, such as ecological niche and resource availability, can act.

However, niche selection or behavioural adaptation alone does not explain why some mosquitoes evolved towards extreme specialism rather than remaining zoophilic. Classic evolutionary theory posits that (host) specialisation is selected for when the fitness benefit of exploiting a particular host outweighs the costs of reduced performance on others (see e.g.[11, 12, 13, 14] (Levins 1962; Futuyma and Moreno 1988; Parvinen and Egas 2004; Forister et al. 2012)).

### 1.2 Constraints and Trade-offs in Mosquito Biting Behaviour

The fitness of a female mosquito depends on two interacting biological systems: (i) host detection, governed by olfactory receptors and neural processing of host-emitted odours [15, 16, 17, 18], and (ii) host exploitation, governed by digestion, nutrient assimilation, and egg production [19]. These systems need not evolve independently. Genetic coupling (e.g. pleiotropy or shared regulation) can impose trade-offs that favour specialisation [20]. To represent these components, we denote by *a*_*g,i*_ the innate attractiveness of mosquito phenotype *g* toward host *i*, and by 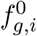 the *basic fitness* (laboratory reproductive payoff) when phenotype *g* feeds exclusively on host *i* (A description of all symbols used in this article can be found in Table 1). If the genetic architecture couples *a*_*g,i*_ and 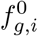, selection on digestive efficiency can “drag” olfactory sensitivity (or vice versa), stabilising specialisation.

**Table 1:**
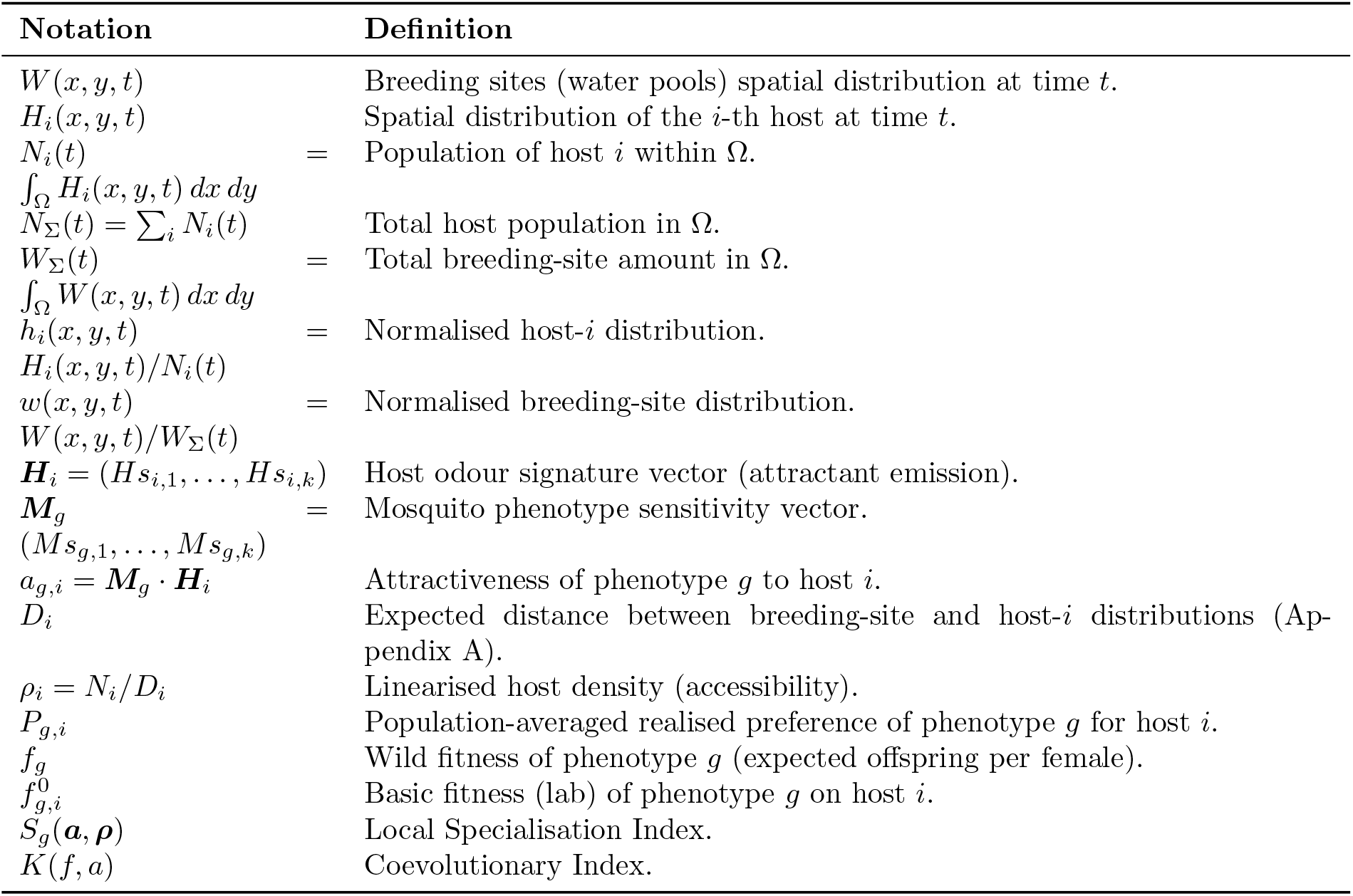
Description of symbols used in the paper.

Regarding host specialisation, this would therefore suggest that anthropophilic mosquitoes feeding on human blood should gain a fitness advantage both over non-specialised mosquitoes as well as over those feeding on off-target hosts. Although empirical studies have repeatedly shown that the blood source can significantly affect mosquito reproductive fitness[21, 22, 23, 24, 25] (Bennett 1970; Richards et al. 2012; Phasomkusolsil et al. 2013; Wekesa et al. 2025; Suresh et al. 2024), this seems to be species-dependent and not always aligned with host preference (as reviewed in [26](de Swart et al. 2023)). In fact, some highly anthropophilic species like *Ae. aegypti* and *An. gambiae* appear to have similar survival and reproduction rates when fed on human or non-human blood, whereas higher reproductive fitness can be found in generalists or zoophilic species, such as *Ae. albopictus, C. quinquefasciatus* or *An. arabiensis*, when fed on preferred host blood. This in turn suggests that the interplay between host availability and realised fitness together influence the evolutionary divergence between specialist and generalist mosquitoes.

### 1.3 A Need for Integrated Spatio-Genetic Modelling

These studies raise key questions: Why does narrow specialisation evolve in some systems but not others? Under what ecological conditions does specialisation outperform opportunism and generalism? How stable is specialisation under environmental change? Addressing these questions requires a framework that integrates genetic constraints, spatial ecology, and temporal variability. A key challenge is that data from laboratory experiments (which quantify intrinsic traits under controlled host access) and field studies (which quantify realised preferences (bite rates) under spatial constraints) are often not directly comparable. The framework below bridges this divide by explicitly incorporating spatial accessibility into the mapping from innate traits to realised preference and fitness.

## 2 Conceptual Eco-Evolutionary Framework

### 2.1 Model overview: tracking realised mosquito fitness in a heterogeneous landscape

This section provides a conceptual overview of the approach, while subsequent sections present the theoretical background and the details of the derivations. The purpose of this model is to track the *realised fitness* of mosquito phenotypes in spatially structured host environments and to determine the ecological and evolutionary conditions under which host specialisation emerges. Specifically, we aim to quantify the expected reproductive output *f*_*g*_ of mosquito phenotype *g* in a landscape containing multiple host species with heterogeneous abundance and spatial configuration. The realised fitness integrates host-seeking behaviour, spatial accessibility, and host-specific nutritional payoffs into a single eco–evolutionary quantity.

At its core, the model links three interacting components:

1. **Host community structure:** Each host species *i* is characterised by (i) its abundance *N*_*i*_, (ii) its spatial distribution relative to mosquito breeding sites, and (iii) its *odour signature*, represented as a vector of emitted chemical compounds. Hosts differ both in abundance and in chemical composition, introducing ecological and sensory heterogeneity into the system.
2. **Mosquito sensory architecture:** Each mosquito phenotype *g* possesses a vector of olfactory sensitivities describing its responsiveness to specific chemical compounds. Mosquitoes differ genetically in their sensitivity profiles, meaning that different phenotypes perceive the same host community differently. The interaction between a host’s odour signature and a mosquito’s sensitivity profile determines the innate attractiveness *a*_*g,i*_.
3. **Host-specific reproductive payoff:** Following a successful blood meal, phenotype *g* obtains a host-dependent reproductive return 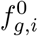, representing physiological efficiency of digestion and egg production under controlled conditions. Hosts differ in nutritional quality, and mosquito phenotypes differ in their ability to exploit those resources.

These three components are embedded in a spatial landscape where breeding sites and host habitats may be separated. Spatial configuration influences effective host accessibility via *the linearised density*, thereby linking ecological geometry to behavioural outcomes.

The model therefore tracks the following causal chain:

Odour emission → Olfactory detection→ Realised host choice → Host-specific fitness→ Total realised fitness.

Diversity is incorporated at two levels. First, multiple host species (*i* = 1, …, *H*) differ in abundance, spatial distribution, chemical signature, and nutritional value. Second, multiple mosquito phenotypes (*g* = 1, …, *G*) differ in sensory sensitivity and physiological efficiency. This bidirectional diversity allows the framework to capture variation both within and between species, making it suitable for analysing polymorphism, trait coupling, and evolutionary divergence.

The quantity we ultimately seek to evaluate is the realised fitness *f*_*g*_, which integrates behavioural allocation across hosts with host-specific reproductive payoff. By expressing realised fitness as a function of host accessibility, sensory architecture, and nutritional efficiency, the model provides a unified structure for analysing when ecological skew becomes evolutionary specialisation.

The following subsections develop each component formally, beginning with host–mosquito sensory interaction and spatial accessibility.

### 2.2 Defining Phenotype, Innate Traits, and Host Landscape

Each mosquito phenotype *g* is characterised by an olfactory sensitivity (to particular chemical compounds) vector ***M***_*g*_ = (*Ms*_*g*,1_, …, *Ms*_*g,k*_), and each host *i* by an odour signature vector ***H***_*i*_ = (*Hs*_*i*,1_, …, *Hs*_*i,k*_). Innate attractiveness is defined as the dot product

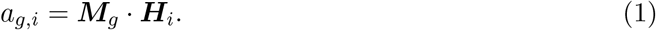

This value describes the innate ability of mosquito brains (i.e., the olfactory signature) to encode host odour features (i.e., the odour signature) for detection and search[27, 28]. Higher co-linearity of the vectors (i.e., the alignment of both signatures) indicates greater innate attractiveness to the host. It should also be noted that, if necessary, the attractiveness vector could potentially incorporate factors unrelated to olfaction, such as temperature, lighting, air movement, and so on.

In turn, the host landscape can be described by a set of spatial distributions*H*_*i*_(*x, y, t*) with a normalised density *h*_*i*_(*x, y, t*) = *H*_*i*_(*x, y, t*)*/N*_*i*_(*t*), where *N*_*i*_ is the total abundance of the *i* − *th* host over region Ω (Table 1).

### 2.3 Spatial Accessibility and Linearised Host Density

Mosquitoes originate from breeding sites with spatial distribution *W* (*x, y, t*) and normalised density *w*(*x, y, t*) = *W* (*x, y, t*)*/W*_Σ_(*t*). The expected distance from breeding sites to host *i* is *D*_*i*_ (formal definition and properties in Appendix A). We define the *linearised host density*

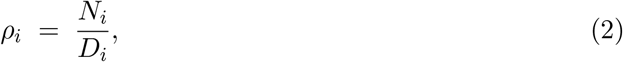

which captures host accessibility as abundance discounted by expected search distance (Figure 1).

**Figure 1:**
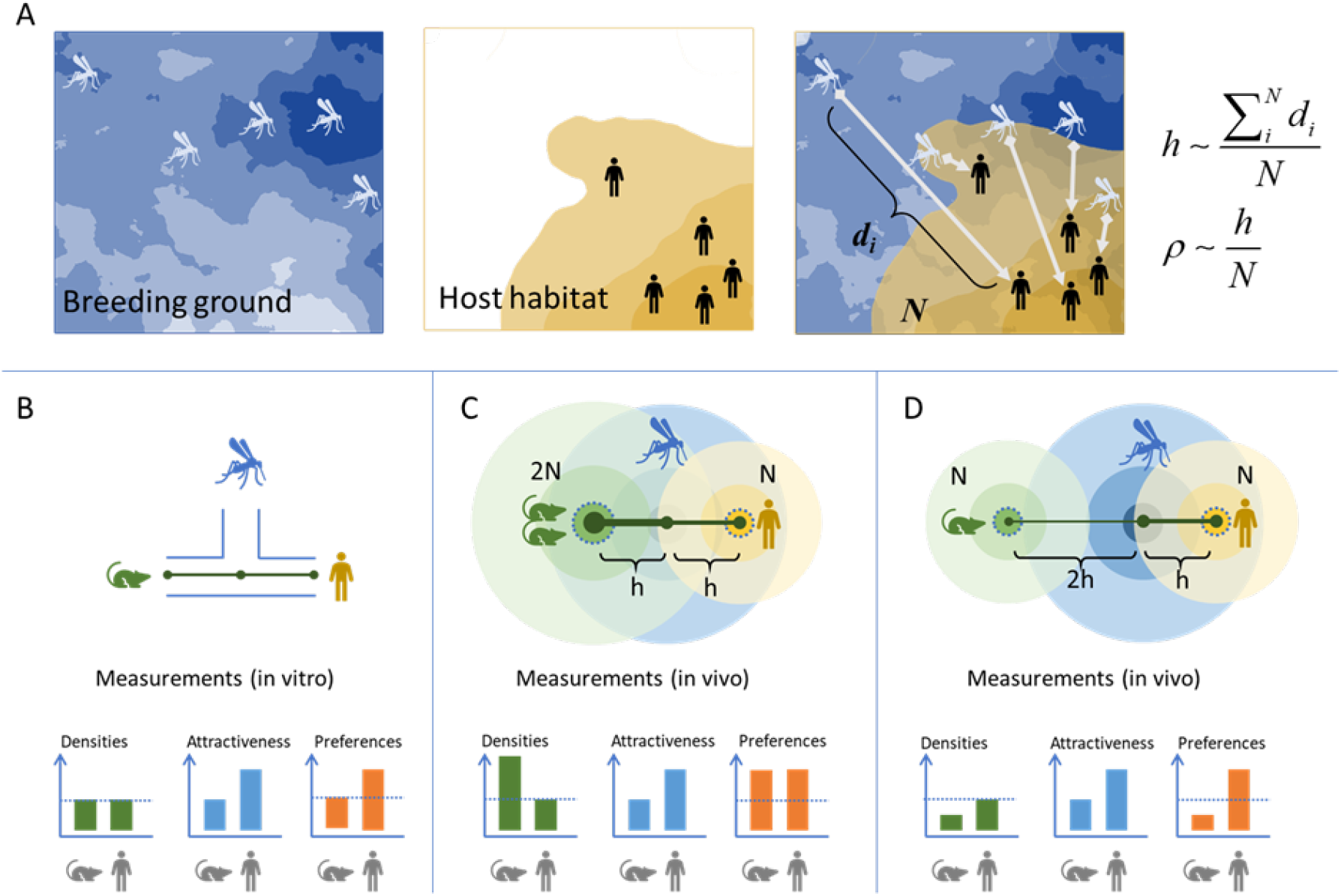
Host-seeking model and linearised host density. (A) Distance metric between breeding-site and host distributions and the concept of linearised host density *ρ*_*i*_ = *N*_*i*_*/D*_*i*_. (B) In vitro choice (standardised laboratory) where measured preferences recover attractiveness. (C)In vivo preferences vs. host abundance at fixed distance. (D) In vivo preferences vs. distance at fixed abundance. (See Appendix A–B for formal definitions and derivations.)

### 2.4 From Innate Traits to Realised Preference

Let *P*_*g,i*_ denote the population-averaged realised preference (fraction of bites) of phenotype *g* on host *i*. Under broad conditions (Appendix B), realised preference depends on the product of innate attractiveness and accessibility:

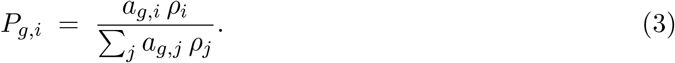

Equation (3) provides the key bridge between in vitro attractiveness and in vivo field-realised preferences, by conditioning the innate attraction on spatial accessibility.

### 2.5 Fitness Architecture: global limitation and local spatial structure

The model tracks the realised fitness *f*_*g*_ of mosquito phenotype *g*, defined as its expected reproductive output under field conditions. Fitness is determined jointly by global resource availability and local spatial structure. Global conditions are represented by the limiting function *L*(*W*_Σ_, *N*_Σ_), where *W*_Σ_ is the total amount of breeding habitat and *N*_Σ_ is the total number of hosts. The relative importance of these resources may differ among ecosystems. For example, mosquito populations in tundra regions may be limited primarily by host scarcity, whereas in some equatorial regions the principal constraint may be the availability of suitable breeding sites. These examples represent contrasting limiting regimes rather than universal properties of those regions.

Local conditions determine which hosts are actually accessible to mosquitoes. They enter the model through the realised preference *P*_*g,i*_, which depends on innate attractiveness and the previously defined linearised host density *ρ*_*i*_. In particular, closer spatial proximity between host-*i* habitats and breeding sites increases host accessibility and therefore increases the contribution of that host to mosquito feeding. Because hosts differ in their basic reproductive payoff 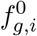, changes in spatial configuration can alter realised mosquito fitness and, consequently, population growth.

The combined fitness architecture is

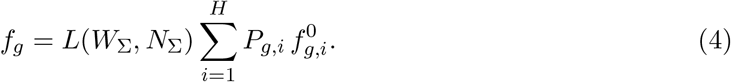

The global function *L* determines the overall ecological capacity for mosquito reproduction, whereas the preference-weighted term describes the local, phenotype-specific return obtained from the accessible host community. Thus, landscapes with the same total numbers of hosts and breeding sites may generate different fitness outcomes when the spatial arrangement of those resources differs. The full matrix formulation and decomposition into environmental, gene–environment, and gene–gene components are given in Appendix C.

### 2.6 Model assumptions and scope

The framework is intentionally minimal and is designed to connect laboratory measurements of mosquito traits with field estimates of host accessibility and realised feeding. Its principal assumptions concern host detection, spatial accessibility, reproductive payoff, and the separation of ecological and evolutionary timescales.

First, the baseline model assumes that the effective signal received from host *i* by phenotype *g* is proportional to *a*_*g,i*_*ρ*_*i*_. Host choice is therefore determined by the relative contribution of each host to the total signal, as expressed in Eq. (3). The accessibility measure *ρ*_*i*_ = *N*_*i*_*/D*_*i*_ is assumed to be finite, with *D*_*i*_ *>* 0. This representation reduces complex spatial structure to an average distance between host and breeding-site distributions; alternative spatial measures may be substituted when more detailed movement or landscape data are available.

Second, realised fitness is represented as a preference-weighted average of host-specific basic fitness, multiplied by the global limiting function *L*(*W*_Σ_, *N*_Σ_) in Eq. (4). The function *L* accounts for constraints imposed by total host availability and total breeding habitat. Within a single environment, it acts as a common multiplier and therefore affects the magnitude of population growth without changing relative fitness rankings among phenotypes.

The model initially treats laboratory estimates of basic fitness as the baseline physiological payoff in the field:

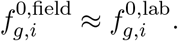

This assumption may require adjustment because field conditions can introduce effects absent from laboratory experiments, including variation in blood-meal size, host defensive behaviour, temperature, infection, and post-feeding survival. Such differences do not alter the model structure. Where necessary, a field-adjusted payoff can be introduced as

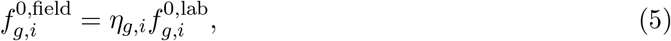

where *η*_*g,i*_ is estimated from field or semi-field data.

Mosquito traits and host characteristics are treated as fixed over a single host-seeking and reproductive period. This separates rapid ecological changes in realised preference, caused by variation in ***ρ***, from slower evolutionary changes in attractiveness and basic fitness. The statistical analysis further assumes that the relevant means, variances, covariances, and higher-order moments are finite, allowing the fitness decomposition presented in Appendix F.

The present framework identifies conditions under which specialisation is favoured, but it does not predict exact evolutionary rates or full mosquito–host population dynamics. Search mortality, unsuccessful host finding, pathogen feedbacks, mutation, migration, and frequency-dependent competition are not modelled explicitly. These processes can be incorporated in future extensions without changing the central distinction between global ecological limitation, local host accessibility, realised preference, and host-specific reproductive performance.

## 3 Quantifying Specialisation

Specialisation cannot be described adequately by the observed proportion of bites on a particular host alone. A mosquito population may feed almost exclusively on one host because it has evolved a narrow intrinsic preference, but the same feeding pattern can arise when that host is simply the only abundant or accessible option. Conversely, a mosquito may possess a genetically constrained host-use system without expressing a strongly specialised diet in every environment. Specialisation must therefore be considered at two related but distinct levels: its *realised ecological expression* and its *underlying evolutionary architecture*.

We introduce two complementary indices to distinguish these levels. The *Local Specialisation Index, S*_*g*_, describes the form of specialisation expressed by mosquito phenotype *g* in a particular host environment. It compares the breadth of the phenotype’s innate attraction profile with the diversity and accessibility of hosts in that environment. The *Coevolutionary Index, K*, describes the structural relationship between the mosquito’s host-detection and host-utilisation systems. It asks whether the ability to find a host and the ability to obtain a reproductive benefit from that host can vary independently or are constrained to evolve together.

The indices are necessary because realised and genetic specialisation need not coincide. A phenotype may have a high local feeding bias but weak coupling between attraction and physiological performance. Such a population is an *ecologically induced* or apparent specialist whose feeding pattern may change rapidly if the host landscape changes. By contrast, a phenotype with both a high Local Specialisation Index and strong sensory–physiological coupling represents a more deeply established specialist whose host association is likely to persist after short-term environmental change.

The two indices therefore answer different questions. The Local Specialisation Index asks, “How specialised is this phenotype under the ecological conditions it currently experiences?” The Coevolutionary Index asks, “How strongly does the underlying trait architecture constrain the phenotype to detect and exploit the same hosts?” Used together, they separate an immediately observed feeding pattern from the evolutionary mechanisms that generate and stabilise it. Their conceptual roles are summarised in Figure 2 and Tables 2 and 3; their formal construction and mathematical properties are given in Appendices D and E.

**Table 2:**
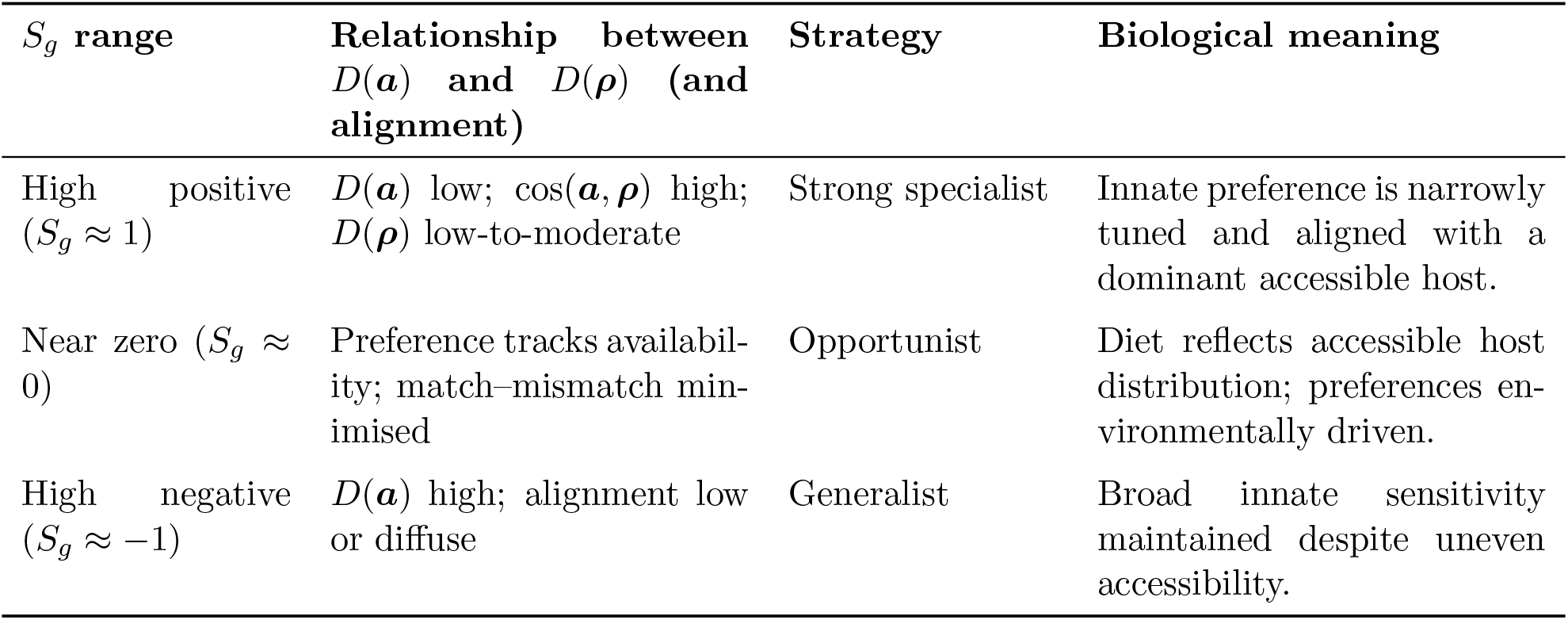
Interpretation of the Local Specialisation Index *S*_*g*_(*a, ρ*).

**Table 3:**
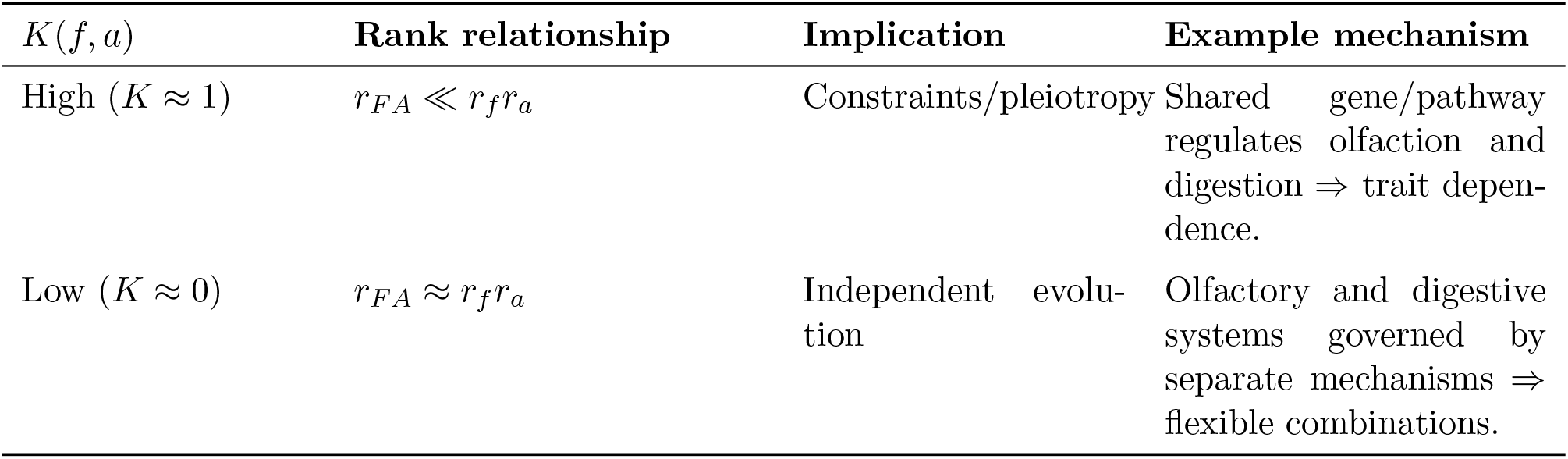
Interpretation of the Coevolutionary Index *K*(*f, a*) based on matrix ranks.

**Figure 2:**
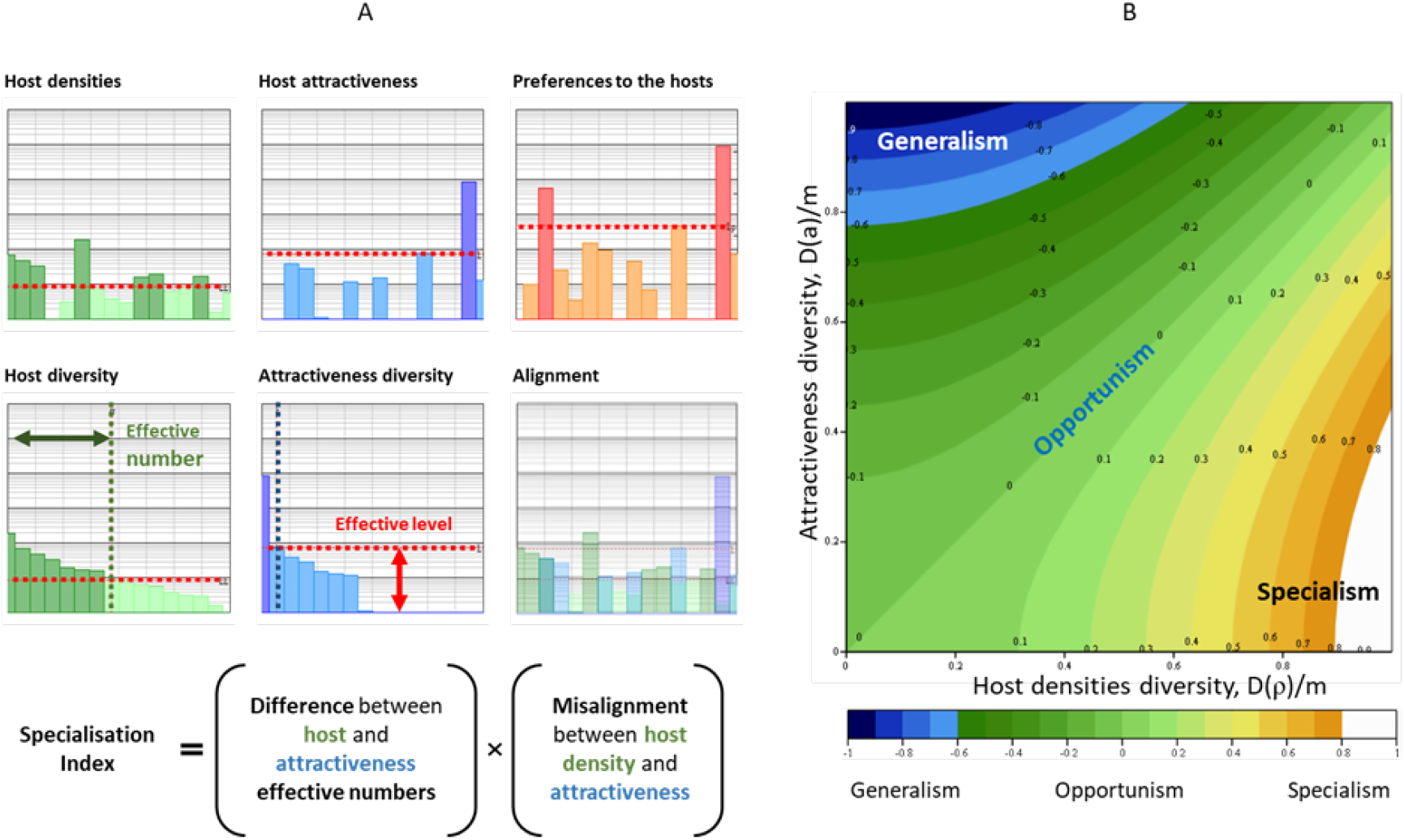
Local Specialisation Index. (A) Construction from host accessibility vector ***ρ***, attractiveness vector ***a***_*g*_, Shannon diversity *D*(·), and cosine alignment. (B) Index values across representative combinations of diversity and alignment. Formal construction and bounds are in Appendix D; interpretation in Table 2.

### 3.1 Local Specialisation Index: realised ecological specialisation

In non-technical terms, the Local Specialisation Index evaluates whether a mosquito’s host-use pattern represents a genuine local preference or merely reflects the choices that the environment makes available. Consider a population in which ninety per cent of observed blood meals come from humans. This pattern does not, by itself, demonstrate intrinsic human specialisation. If humans account for nearly all accessible hosts, a broadly feeding mosquito could produce the same observation. Stronger evidence of specialisation would be obtained if several hosts were readily accessible, but the mosquito remained narrowly attracted to humans. The index is intended to distinguish these situations.

The calculation uses two vectors. The attractiveness vector ***a***_*g*_ = (*a*_*g*,1_, …, *a*_*g,m*_) describes the innate responsiveness of phenotype *g* to the available host species. The accessibility vector ***ρ*** = (*ρ*_1_, …, *ρ*_*m*_) describes the same host community after the abundance and spatial proximity to the breeding sites have been taken into account. The index compares the effective diversity of these vectors and measures their alignment. Thus, it incorporates both the *breadth* of innate host detection and the degree to which that detection is directed toward hosts that can actually be encountered. Define the effective diversity based on Shannon-entropy *D*(·) and cosine alignment cos(***a***_*g*_, ***ρ***). The Local Specialisation Index is

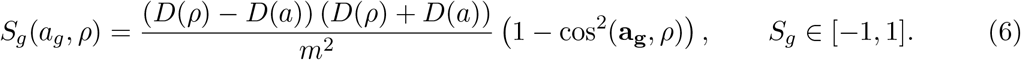

The complete construction, including normalisation and limiting cases, is presented in Appendix D.

The sign and magnitude of *S*_*g*_ describe three broad ecological strategies. A positive value indicates that the phenotype’s innate attraction profile is narrower than the accessible host community and is directed toward hosts present in that community. The phenotype therefore expresses local specialisation: it selects a restricted subset even though a broader set of hosts is potentially available. A value near zero indicates opportunism. In this case, the breadth of attraction approximately matches the breadth of local host accessibility, so feeding is largely determined by the environment rather than by a strong intrinsic restriction. A negative value indicates a generalist tendency, in which the mosquito retains a broader attraction profile than the host diversity currently available to it.

It is important to distinguish strictly between opportunism and generalism, although the two terms are often used interchangeably in the ecological literature. In the present framework, *opportunism* describes a realised feeding pattern that tracks the local availability of hosts, whereas *generalism* describes an intrinsically broad host-use phenotype capable of detecting and exploiting several host types. Generalism is therefore not treated simply as the absence of specialisation, but as a multi-host form of specialisation: the phenotype is adapted to a broad set of hosts rather than narrowly adapted to a single host.

The term *local* is important because *S*_*g*_ is explicitly environment-dependent. The same phenotype can have different values of *S*_*g*_ in different landscapes or seasons even when its innate attractiveness vector remains unchanged. For example, a phenotype may appear opportunistic in a host-poor dry-season environment but exhibit a clear preference when several hosts become accessible during the wet season. Changes in *S*_*g*_ over time do not necessarily imply genetic evolution; they may instead represent the changing ecological expression of a fixed sensory phenotype.

The index is also distinct from realised preference *P*_*g,i*_. Equation (3) reports the fraction of feeding directed toward each host, whereas *S*_*g*_ helps interpret whether that pattern reflects intrinsic narrowness, environmental forcing, or a match between the two. Empirically, estimating *S*_*g*_ therefore requires more information than blood-meal proportions alone. Innate attraction can be estimated from controlled host-choice or olfactometer experiments, while *ρ*_*i*_ can be estimated from host abundance and the spatial relationship between host and breeding-site distributions. Figure 2 illustrates the construction of the index, and Table 2 summarises its biological interpretation.

### 3.2 Coevolutionary Index: genetic and functional specialisation

The Coevolutionary Index addresses a different question. In non-technical terms, it asks whether mosquitoes that are good at finding a particular host are also constrained to be good at converting that host’s blood into offspring. Detection and utilisation are biologically distinct processes: one is governed primarily by sensory and neural systems, while the other depends on digestion, metabolism, and reproduction. If these systems can evolve independently, a mosquito can retain broad host detection while becoming physiologically efficient on one host, or it can change its host preference without a corresponding change in blood-meal performance. If the systems are tightly coupled, however, adaptation in one component restricts the possible states of the other and makes narrow specialisation more likely to persist.

To represent this distinction, we consider the variation between multiple mosquito phenotypes and multiple host species. Let ***A*** = [*a*_*g,i*_] be the phenotype-by-host attractiveness matrix and let 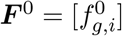 be the corresponding matrix of host-specific basic fitness. Their elementwise, or Hadamard, product is ***F A*** = ***F***^***0***^◦ ***A***. This interaction matrix records the combined contribution of detection and physiological performance for every phenotype–host combination. The ranks of these matrices describe the number of independent dimensions required to represent variation in the corresponding traits. We write *r*_*a*_ = rank(***A***), *r*_*f*_ = rank(***F*** ^0^), *r*_*F A*_ = rank(***F A***).

The Coevolutionary Index is defined as

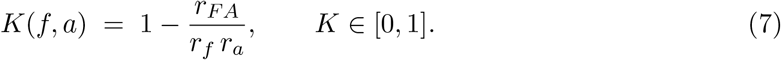

The rank inequality for a Hadamard product ensures that *r*_*F A*_ ≤ *r*_*f*_ *r*_*a*_. The derivation, admissible range, dimensional constraints, and numerical implementation are described in Appendix E.

A relatively high value of *K* indicates that the combined attractiveness–fitness matrix has substantially fewer independent dimensions than would be expected from the separate trait matrices. Biologically, this pattern is consistent with a restricted set of viable detection–utilisation combinations. Such restriction could arise from pleiotropy, genetic linkage, shared regulation, developmental integration, or another mechanism that causes sensory and physiological traits to vary together. In this regime, adaptation toward a particular host in one system tends to be accompanied by adaptation in the other, stabilising host association and reducing the evolutionary accessibility of alternative strategies.

A relatively low value of *K* indicates that the combined interaction retains more of the dimensional complexity present in the separate trait systems. Detection and utilisation are then comparatively free to vary in different combinations. This flexibility can support generalism, host switching, behavioural plasticity, or mixed strategies. It also means that a high Local Specialisation Index need not indicate a genetically entrenched state: feeding may remain strongly biased only while the current ecological conditions persist.

The Coevolutionary Index should be interpreted as a structural indicator of trait coupling rather than as direct proof of a particular genetic mechanism. A high value can identify a pattern consistent with pleiotropy or shared regulation, but genomic, breeding, or gene-expression studies are required to determine its molecular cause. In empirical applications, ***A*** and ***F*** ^0^ should be estimated across several phenotypes and hosts. Because exact matrix rank is sensitive to measurement error, Appendix E also describes the use of singular-value thresholds and null or permutation models for estimating effective rank and determining whether the observed coupling exceeds that expected from sampling noise and matrix dimensions.

Joint interpretation of *S*_*g*_ and *K* provides a more complete classification of specialisation. High *S*_*g*_ together with high *K* indicates locally expressed and structurally stabilised specialisation. High *S*_*g*_ with low *K* indicates a feeding bias that is likely to be environmentally induced or evolutionarily reversible. Low or near-zero *S*_*g*_ with high *K* indicates that a constrained trait architecture exists but is not strongly expressed in the current
current environment. Finally, low *S*_*g*_ and low *K* are consistent with a flexible generalist or opportunistic strategy. The two indices therefore distinguish what mosquitoes currently do from the degree to which their biological architecture commits them to continue doing it.

## 4 Results

### 4.1 Static Environments: fitness decomposition and conditions for specialisation

We first consider a static environment in which total host abundance, breeding-site availability, and the spatial distributions of hosts and breeding sites remain constant. Under these conditions, the global limiting function *L*(*W*_Σ_, *N*_Σ_) is constant and sets only the overall scale of mosquito population growth. Differences in realised fitness among mosquito phenotypes are therefore determined by local factors: host accessibility *ρ*_*i*_, innate attractiveness *a*_*g,i*_, and host-specific basic fitness 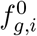.

As shown in Appendix F, Equation (4) can be decomposed into five contributions. The first is determined by the mean values of accessibility, attractiveness, and basic fitness. The next three are covariance terms describing pairwise alignment between attractiveness and accessibility, basic fitness and accessibility, and attractiveness and basic fitness. The final term is a co-skewness term describing a higher-order association among all three variables. This decomposition separates broad improvement across hosts from host-specific alignment.

Realised fitness can consequently increase through two principal routes. The first is an increase in mean performance. Selection may favour mosquitoes with greater average attractiveness or greater average basic fitness across the host community. This pathway improves performance broadly and therefore tends to support generalism. Mean accessibility is an environmental property rather than a mosquito trait, but a landscape with higher average accessibility can likewise raise the overall opportunity for successful host use.

The second route is an increase in the positive covariance terms. A positive association between attractiveness and accessibility means that mosquitoes are especially responsive to hosts that are locally available. A positive association between basic fitness and accessibility means that the most accessible hosts also provide relatively high reproductive returns. A positive association between attractiveness and basic fitness means that mosquitoes preferentially detect the hosts on which they perform best physiologically. These covariance terms generate the host-specific advantages that can favour specialisation. In particular ecological configurations, the co-skewness term can strengthen or weaken this effect when the same restricted host subset is simultaneously accessible, attractive, and reproductively profitable, even when the pairwise correlations alone do not fully describe the pattern.

The relative importance of these terms depends on the structure of the host landscape. Here, a *poor landscape* is one in which only one or a few host species are effectively available to mosquitoes, so accessibility is strongly uneven across hosts. A *rich landscape* contains many host species with approximately equal accessibility. These terms refer to the diversity and evenness of host *availability*, not to total host abundance: a landscape may contain many individual hosts and still be locally poor if almost all accessible hosts belong to one species.

By considering landscape richness together with the magnitude and direction of the correlations among *ρ, a*, and *f* ^0^, the decomposed fitness expression identifies several possible evolutionary pathways. In poor landscapes, variation in accessibility tends to dominate, and specialisation is directed toward the small number of hosts that can be reached. In rich landscapes, spatial constraints are weaker, and the outcome depends more strongly on variation in attractiveness and basic fitness. Selection may then favour physiological specialisation on high-quality hosts, sensory specialisation on readily detectable hosts, or coordinated specialisation when detection and utilisation traits are aligned. These pathways are illustrated in Figure 3 and summarised in Table 4.

**Table 4:** Evolutionary pathways in static environments (synthesis of Figure 3).

| Variation ratio | Dominant selection pressure | Likely outcome | Context |
| --- | --- | --- | --- |
| $cv(\rho) \gg cv(a), cv(f^0)$ | Host accessibility $\rho$ | Narrow specialisation on accessible hosts | “Poor” environments (few accessible hosts) |
| $cv(\rho) \ll cv(a), cv(f^0)$ | Co-specialisation (interaction) | Multiple co-specialised phenotypes | “Rich” environments (many accessible hosts) |
| $cv(f^0) \gg cv(a)$ | Basic fitness variation | Physiological specialisation on high-quality blood | Selection targets utilisation |
| $cv(a) \gg cv(f^0)$ | Attractiveness variation | Sensory specialisation on detectable hosts | Selection targets perception |

**Figure 3:**
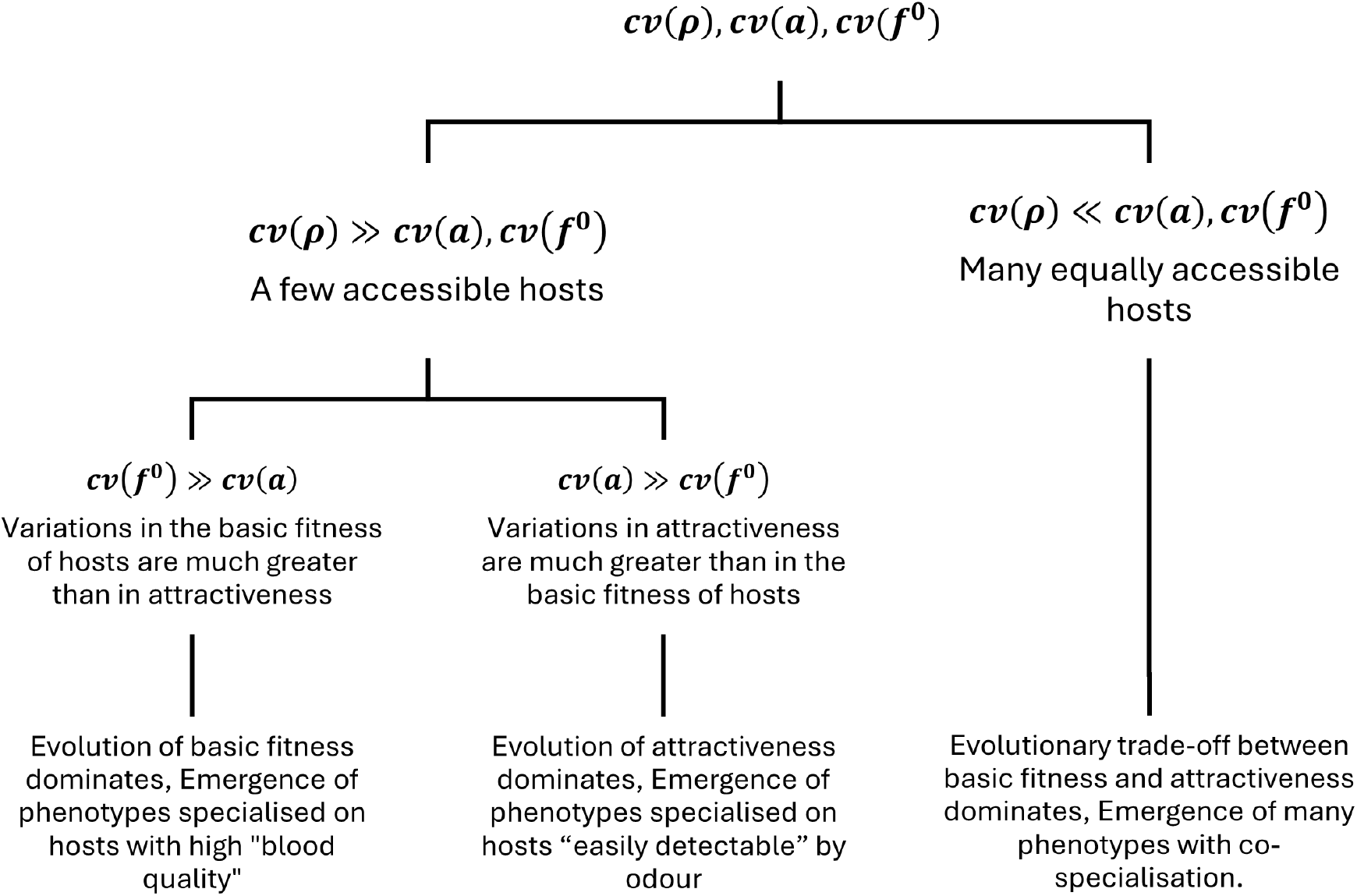
Evolutionary pathways as a function of variability ratios. Branching regimes predicted by relative magnitudes of cv(*ρ*), cv(*a*), and cv(*f* ^0^). The classification corresponds to leading terms in the fitness expansion (Appendix F) and is summarised in Table 4.

The principal conclusion is that the evolution of specialisation requires persistent correlations. These may occur between mosquito traits and the environment, such as correlations of attractiveness or basic fitness with host accessibility, or between mosquito traits themselves, particularly between attractiveness and basic fitness. Trait–environment correlations can produce strong but locally contingent specialisation that may weaken when the landscape changes. By contrast, correlation between host detection and host utilisation indicates co-adaptation within the mosquito phenotype and is therefore expected to produce a deeper, more stable, and potentially less reversible form of specialisation. This distinction links the fitness decomposition to the complementary roles of the Local Specialisation Index and the Coevolutionary Index.

### 4.2 Worked Ecological Scenarios: Predictable Regimes of Specialisation

We now interpret the model through ecological scenarios described by the relative variability of accessibility and intrinsic traits.

#### Scenario I: Resource-poor landscapes (forced specialisation by accessibility)

When *ρ* is highly uneven across hosts (large cv(*ρ*)), realised preference (3) concentrates on the few accessible hosts even if innate attraction is broad. Selection therefore favours improving performance on those hosts, producing rapid specialisation. Apparent specialisation can be strong in this regime even before substantial genetic adaptation occurs. This regime is expected in landscapes where breeding sites are spatially concentrated near a subset of hosts (e.g. urban container habitats near humans; stable wetlands near waterfowl).Figure 4(A).

**Figure 4:**
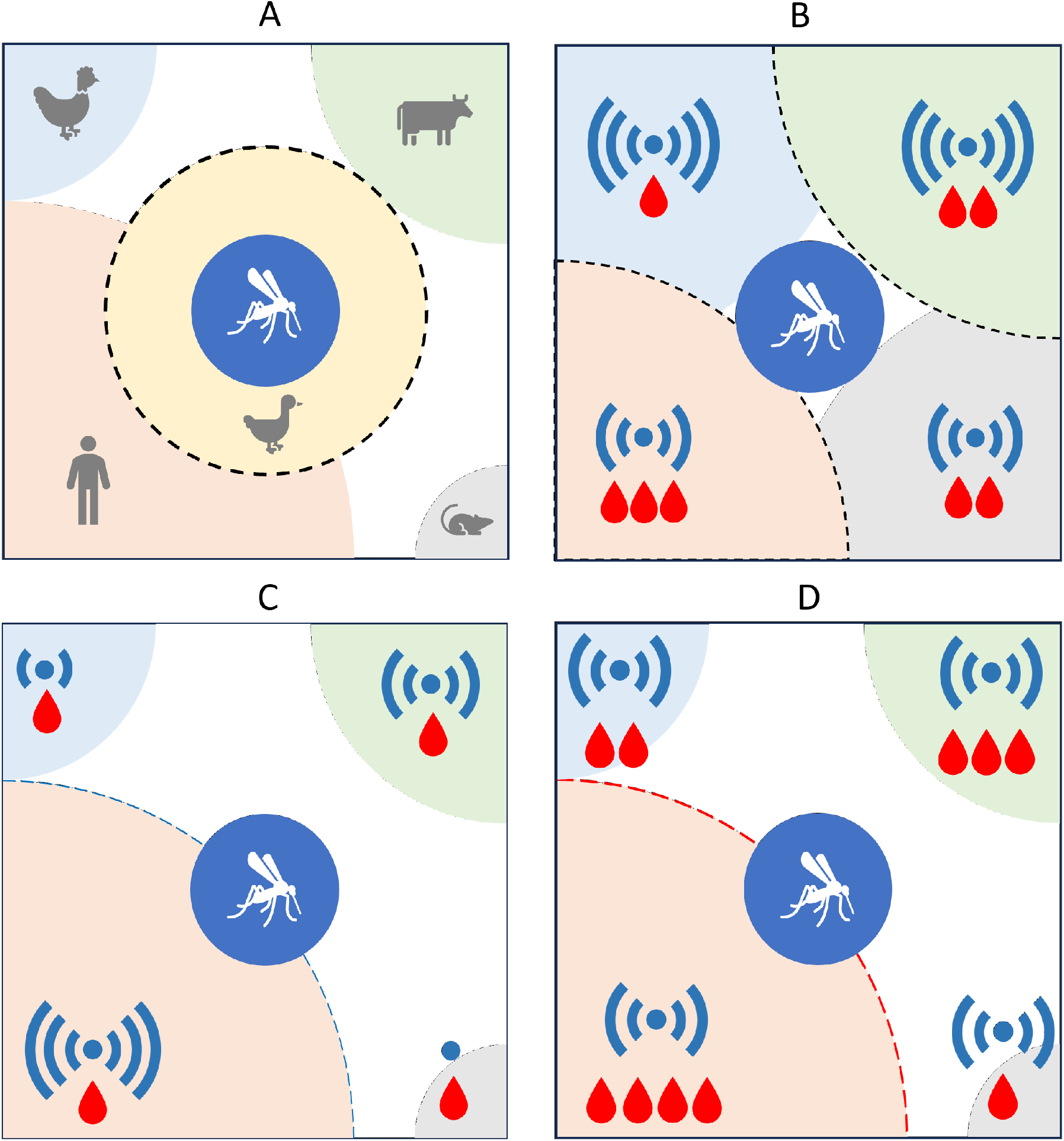
Mosquito specialisation scenarios. Each panel shows a hypothetical landscape with the mosquito habitat (blue circle) at the centre and host habitats (coloured sectors) at the corners of the square. (A) **Scenario I**: A single host completely surrounds the mosquito habitat. In this case, the specialisation index is always high, regardless of the coevolutionary index. (B) **Scenario III**: All hosts are equally accessible; here, specialisation depends on the correlation between attractiveness and basic fitness (i.e., mosquitoes will specialise on the host with the optimal combination of both traits). (C) **Scenario IIa**: There is a correlation between host accessibility and attractiveness; in this case, specialisation is driven by attractiveness. (D) **Scenario IIb**: There is a correlation between accessibility and basic fitness; in this case, specialisation is driven by basic fitness.

#### Scenario II: Resource-rich landscapes (trait-driven specialisation)

When hosts are similarly accessible (small cv(*ρ*)), spatial constraints do not dominate preference. Selection shifts to intrinsic traits: if cv(*f* ^0^) ≫ cv(*a*), physiological specialisation on high-quality blood dominates (Scenario IIb, Figure 4(D)); if cv(*a*) ≫ cv(*f* ^0^), sensory specialisation on highly detectable hosts dominates (Scenario IIa, Figure 4(C)). Thus the dominant type of specialisation is predicted by the leading axis of trait variance.

#### Scenario III: Genetic coupling and co-specialisation

When both *a* and *f* ^0^ vary and are genetically coupled (high *K*), the interaction structure ***F A*** becomes low-rank relative to *r*_*f*_ *r*_*a*_. The fitness landscape fragments into multiple constrained optima, favouring the emergence of multiple co-specialised strategies (host-associated phenotypes) rather than a single global specialist.Figure 4(B).

An extreme limiting case of Scenario III can also be envisaged in which accessibility, attractiveness, and basic fitness are jointly aligned: the most nutritionally valuable hosts are simultaneously the easiest to detect and the most accessible. Under these conditions, the three-way co-skewness term in the fitness decomposition may become dominant, reinforcing particularly strong specialisation. Although this configuration is mathematically possible in a static environment, such alignment is less likely to persist under seasonal variation, because host accessibility, detectability, and nutritional value need not covary through time. We therefore treat it as an extreme extension of Scenario III rather than as a separate evolutionary pathway.

### 4.3 Seasonality as an Evolutionary Filter

Seasonality affects both *N*_*i*_(*t*) and the breeding site distribution *W*(*x, y, t*), thereby altering *ρ*_*i*_(*t*) and potentially changing the sign and magnitude of trait correlations. If seasonal changes are proportional and preserve correlation signs, they can be evolutionarily neutral in the sense of not changing the direction of selection.

By contrast, seasonal processes that reverse correlation signs (e.g. migrations, breeding habitat drying, host relocation) can generate strong evolutionary filtering. Two pathways follow: (i) *over-adaptation to the limiting season* (e.g. dry season) and (ii) *specialisation on seasonal refuges* (hosts whose accessibility remains stable across seasons). The emergence of narrow specialisation is therefore predicted to be concentrated in “hot spot” ecosystems with temporally stable spatial correlation between breeding sites and host habitats (e.g. urban and marsh systems).

### 4.4 Simulated Transition to Host Specialisation

To illustrate the eco-evolutionary transition predicted by the framework, we simulated a gradual change in accessibility mimicking urbanisation or increasing the severity of the dry-season: the accessibility to host 2 *ρ*_2_(*t*) increases while the accessibility to host 1 *ρ*_1_(*t*) decreases (simulation methods and update rules in Appendix F, Section F.6). Initially, accessibility is balanced, and the population exhibits opportunistic feeding (*S*_*g*_ ≈ 0), with realised preference tracking availability. As accessibility shifts, realised preference rapidly moves toward host 2 (apparent specialisation), producing a sharp rise in the local specialisation index *S*_*g*_. Under moderate to strong genetic coupling (high *K*), the interaction between *a* and *f* ^0^ stabilises: *K* rises more slowly, indicating longer-term co-adaptation and persistence of host specialisation even if ecological forcing weakens. Figure 5 summarises these dynamics in a single transition vignette.

**Figure 5:**
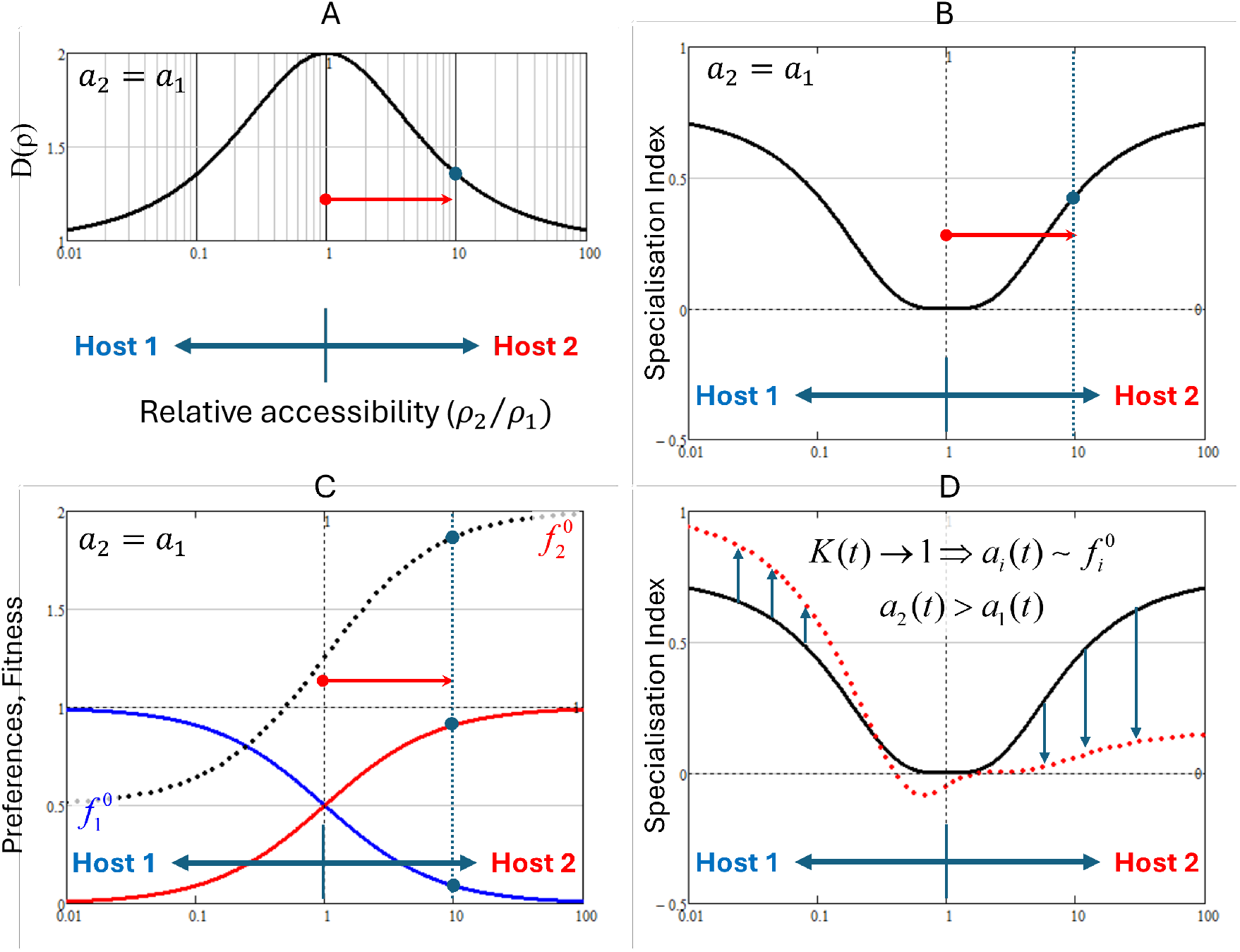
Simulated eco-evolutionary transition to host specialisation. (A) Gradual environmental shift in relative host accessibility (*ρ*_2_*/ρ*_1_ *>* 1,*ρ*_1_(*t*) ↓, *ρ*_2_(*t*) ↑) is shown by the red arrow. The top graph shows the effective host number *D*(*ρ*) as a function of relative accessibility. (B) Local Specialisation Index *S*_*g*_ responds quickly to ecological forcing. The graph shows the specialisation index as a function of relative accessibility, given equal host attractiveness(*a*_2_ = *a*_1_). (C) Realised preference shifts rapidly toward host 2 (apparent specialisation). The graph shows the changes in preferences (blue line - host 1, red line - host 2) and the realised fitness for these preferences. (D) Coevolutionary Index *K* rises more slowly, indicating longer-term genetic coupling and stabilisation. The graph shows the change in the specialisation index with changes in the attractiveness of the hosts (*a*_2_ *> a*_1_). Simulation update rules and parameterisation are provided in Appendix F (Section F.6).

## 5 Discussion

### 5.1 Ecology restricts, genetics stabilise: a unified view of host specialisation

The framework developed here provides a unifying interpretation of mosquito host specialisation as an eco–evolutionary process structured by two interacting filters: ecological gating and genetic stabilisation. The ecological gate is determined by spatial accessibility through the linearised host density *ρ*_*i*_ (Eq. (2)). In contrast, the genetic filter is determined by the coupling between detection and exploitation traits, captured by the Coevolutionary Index *K* (Eq. (7)). Together, these determine whether apparent feeding bias remains transient and context-dependent, or becomes evolutionarily entrenched.

Equation (4) shows that realised fitness depends on the interaction of innate attractiveness *a*_*g,i*_, accessibility *ρ*_*i*_, and host-specific payoff 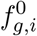. Crucially, specialisation is not favoured merely because one host is abundant or nutritionally superior; it requires persistent alignment between trait structure and environmental structure (Appendix F). When this alignment exists, covariance terms in the fitness expansion (Eq. F1–F2) generate directional selection toward narrow host use. When alignment is absent, selection favours generalist optimisation.

This leads to a central conclusion: *host specialisation is not an intrinsic species property but an emergent property of trait–environment correlation structure*. This perspective reconciles long-standing debates between intrinsic-preference and opportunistic-feeding interpretations in vector ecology.

### 5.2 Reinterpreting apparent specialisation in field studies

Field observations often classify mosquito species as specialists or generalists based on blood-meal analysis. However, Eq. (3) implies that realised preference can be strongly skewed purely by accessibility effects. The Local Specialisation Index *S*_*g*_ (Eq. (6)) was designed precisely to distinguish between intrinsic narrowness of sensory architecture and environmental forcing.

A high proportion of bites on a single host does not necessarily imply evolutionary specialisation. If host diversity is low (small *D*(***ρ***)) and attractiveness diversity is high, *S*_*g*_ may remain near zero or negative despite extreme bite skew. Conversely, modest bite skew may reflect strong intrinsic bias if the environment is diverse.

This distinction is particularly important for interpreting host shifts under anthropogenic change. Urbanisation often increases *ρ*_human_ by simultaneously increasing host abundance and reducing distance between breeding sites and humans. Under such conditions, generalist populations may appear to specialise behaviourally before any genetic shift occurs[29]. Longitudinal monitoring of both *S*_*g*_ and *K* would allow separation of ecological forcing from genetic canalisation.

### 5.3 Seasonality as a directional evolutionary filter

The role of seasonality emerges naturally from the structure of Eq. (4). Seasonal variation modifies *ρ*_*i*_(*t*) and therefore the trait–environment covariance terms that drive selection. If seasonal fluctuations preserve the sign of Cov(*aρ, f* ^0^) across time, selection direction remains stable. If fluctuations reverse that sign, directional selection is weakened or oscillatory. Two distinct evolutionary regimes follow: **Stable seasonal correlation:** When host accessibility patterns repeat predictably (e.g. human dominance in urban dry seasons), specialisation accumulates across generations. Such environments function as evolutionary ratchets. **Unstable correlation:** When seasonal changes invert accessibility patterns (e.g. migratory hosts or temporary livestock introduction), selection alternates direction, favouring flexible or generalist strategies unless strong genetic coupling (*K* → 1) constrains plasticity.

This framework predicts that the most extreme specialisations should be observed in ecosystems where spatial correlations between breeding sites and host habitats remain temporally stable, such as permanent marshes or dense urban settlements.

### 5.4 Genetic coupling and the irreversibility of specialisation

The Coevolutionary Index *K* formalises an often implicit concept: specialisation becomes evolutionarily stable when detection and exploitation traits collapse onto a shared axis of variation. When *K* is high, the effective dimensionality of trait space (Appendix E) shrinks, restricting possible trait combinations. In such systems, improvement in host detection inherently improves host exploitation, reinforcing directional selection.

This provides a mechanistic explanation for why some mosquito lineages appear locked into narrow host use even when alternative hosts become available. Once the genetic mechanisms governing olfaction and digestion become tightly coupled (i.e., a rank collapse occurs in **F**^0^ ◦ **A**), reverting to opportunism requires re-expansion of trait dimensionality — an unlikely evolutionary reversal.

By contrast, low *K* implies that sensory and physiological systems can adjust independently. In such populations, behavioural plasticity or partial shifts are more feasible, and specialisation may remain reversible.

The simulated transition (Figure 5) illustrates a two-phase process: **Rapid ecological skew:** Changes in *ρ*_*i*_ immediately alter realised preference via Eq. (3), producing high *S*_*g*_ even without genetic change. **Gradual genetic stabilisation:** Under selection gradients derived from Eq. F1, trait evolution increases alignment between *a* and *f* ^0^, raising *K* and embedding specialisation in genetic architecture.

This distinction mirrors empirical observations in invasion biology and urban adaptation, where behavioural shifts precede detectable genomic differentiation. The model therefore offers a quantitative scaffold for interpreting eco-evolutionary transitions during environmental change.

### 5.5 Implications for vector-borne disease dynamics and vector control

Host specialisation alters disease transmission in multiple ways. Narrow specialisation increases vector–host contact frequency within a specific host class, potentially amplifying transmission when that host is a competent reservoir. Conversely, generalist feeding can dilute transmission by distributing bites across non-competent hosts.

Because specialisation emerges most strongly in accessibility-stable environments, disease risk may intensify in habitats where breeding sites and competent hosts remain spatially correlated over time. Urban consolidation of breeding habitats near human populations may therefore not only increase vector density but also favour evolutionary specialisation on humans.

The framework predicts that interventions disrupting spatial correlation — such as separating breeding sites from dominant hosts — may reduce not only immediate biting rates but also long-term evolutionary pressure toward specialisation.

The eco–evolutionary framework developed in this study also offers practical insights for vector management and disease control strategies. By explicitly linking mosquito behaviour and evolutionary trajectories to spatial host accessibility through the linearised host density *ρ*_*i*_ = *N*_*i*_*/D*_*i*_, the model highlights how environmental structure can indirectly shape vector host preference and therefore pathogen transmission risk.

A key implication is that control strategies targeting only mosquito population size may overlook an equally important driver of disease transmission: the spatial correlation between breeding sites and preferred hosts. When breeding sites are located close to a dominant host species, the accessibility parameter *ρ*_*i*_ becomes large, which increases the probability of repeated feeding on that host (Eq. (3)). Such spatial alignment can promote both behavioural skew in feeding patterns (high *S*_*g*_) and eventual evolutionary stabilisation of host specialisation (high *K*). In this context, vector control efforts that merely reduce mosquito abundance without disrupting this spatial alignment may fail to prevent the emergence of specialised vector populations.

The framework therefore suggests that vector management should focus on *decoupling breeding habitats from dominant hosts*. For example, urban mosquito control programs often concentrate on eliminating standing water containers. In addition to reducing mosquito density, these measures may also decrease host accessibility *ρ*_*i*_ for humans by increasing the average distance between breeding sites and human dwellings. By lowering *ρ*_human_, such interventions reduce the ecological advantage of human-specialist strategies and may favour a shift toward broader host use or opportunistic feeding.

Similarly, landscape-level interventions that redistribute or buffer host populations may alter the correlation structure between mosquito traits and host accessibility. Zooprophylactic strategies—introducing alternative animal hosts to divert mosquito bites away from humans[30]—can be interpreted in this framework as attempts to increase host diversity *D*(*ρ*) and thereby reduce the effective alignment between mosquito attractiveness vectors and human accessibility. When successful, this strategy decreases *S*_*g*_ for human hosts and may weaken selection pressures favouring human specialisation.

Another important implication concerns environmental stability. The model predicts that narrow host specialisation is most likely to emerge in ecosystems where the spatial relationship between hosts and breeding sites remains stable over long periods. Urban environments, where water storage containers and human hosts are consistently co-located, represent such conditions[31]. In contrast, landscapes with fluctuating host distributions or temporary breeding habitats—such as agricultural systems or floodplains— may maintain mosquito populations in a more generalist or opportunistic state. Consequently, environmental variability itself may function as a natural buffer against the evolution of extreme host specialisation.

Finally, the proposed indices *S*_*g*_ and *K*(*f, a*) may serve as practical diagnostic tools in surveillance programs. Monitoring realised feeding patterns together with ecological accessibility could allow estimation of the Local Specialisation Index *S*_*g*_, indicating whether observed host bias is primarily ecological or genetically entrenched. Long-term genomic or physiological studies may then estimate *K*(*f, a*) to determine whether host preference has become evolutionarily constrained. Such metrics could provide early warning indicators of emerging human-specialist mosquito populations and guide targeted intervention strategies.

Taken together, these insights suggest that effective vector control may require not only reducing mosquito abundance but also reshaping the spatial structure of host–vector interactions. By altering the ecological parameters that determine host accessibility and trait–environment correlations, management interventions may influence both the short-term behavioural ecology and the long-term evolutionary trajectories of mosquito vectors.

### 5.6 Limitations and modelling extensions

Several simplifying assumptions warrant further investigation: **Additive fitness structure:** Equation (4) assumes additive host payoffs. Nonlinear digestion efficiencies or density-dependent host depletion could introduce frequency-dependent selection. **Static trait architecture:** Genetic variation is treated abstractly through rank measures. Explicit quantitative genetic models would allow investigation of evolutionary rates and polymorphism stability. **Absence of pathogen feedback:** Pathogen-induced behavioural changes or fitness costs are not incorporated. Coupling this framework with epidemiological dynamics may reveal feedback loops between specialisation and pathogen persistence. **Epigenetic modulation:** Non-genetic inheritance may transiently alter effective trait structure, influencing early-stage specialisation before genomic fixation[32, 33, 34].

Future extensions could integrate density-dependent mosquito population dynamics, host demography, and stochastic seasonal forcing to explore conditions under which polymorphism, branching, or collapse of specialisation occurs.

## 6 Conclusion

By formally linking spatial ecological constraints (linearised host density *ρ*) to innate genetic potential (*a* and *f* ^0^), this framework shows that mosquito host specialisation is an environmentally gated outcome that is further shaped by genetic coupling and seasonality. The indices *S*_*g*_ and *K* provide practical, testable metrics to disentangle ecological opportunism from evolutionary specialisation, offering a foundation for empirical validation and predictive risk assessment.

Although developed for mosquitoes, the structure of the model is general. Any consumer species facing heterogeneous resource landscapes and trait-mediated detection–exploitation trade-offs may exhibit analogous eco–evolutionary gating. The separation of ecological skew (*S*_*g*_) from genetic coupling (*K*) provides a template for distinguishing plastic niche compression from true adaptive specialisation in other systems.

By formalising host specialisation as a covariance-driven, dimension-reducing process in trait–environment space, this framework contributes to a broader synthesis linking spatial ecology, functional trait architecture, and evolutionary constraint theory.

## A Appendix A

Spatial Distance Metric and Linearised Host Density

### A.1 Expected distance between breeding sites and host habitats

Let Ω ⊂ ℝ ^2^ denote the landscape region. Let *w*(***x***) be the normalised breeding-site density and *h*_*i*_(***y***) the normalised host-*i* density. For two points ***x*** = (*x*_*A*_, *y*_*A*_) and ***y*** = (*x*_*B*_, *y*_*B*_) drawn independently from *w* and *h*_*i*_, define Euclidean distance

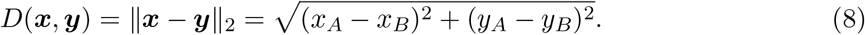

The expected distance from breeding sites to host *i* is

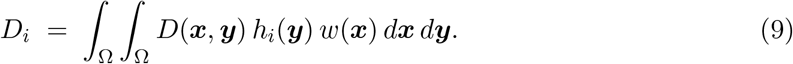

Existence: if Ω is bounded and *w, h*_*i*_ are integrable densities, then *D*_*i*_ *<* ∞.

### A.2 Linearised host density

Define linearised host density

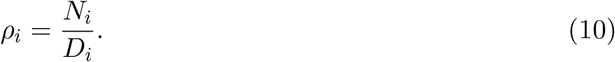

Interpretation: *ρ*_*i*_ increases with abundance and decreases with spatial separation between breeding sites and host habitat; it is a scalar proxy for accessibility under the assumption that encounter probability decreases with distance (formalised in Appendix B).

## B Appendix B Derivation of Realised Preference

### B.1 Effective signal

Let *c*_*g,i*_ be the effective attractant signal from host *i* perceived by phenotype *g*. Assume *c*_*g,i*_ is proportional to (i) innate attractiveness *a*_*g,i*_, (ii) host abundance *N*_*i*_, and decreases with expected distance *D*_*i*_:

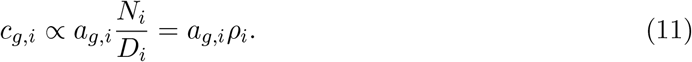

Let *C*_*g*_ = Σ_*j*_ *c*_*g,j*_ be total signal. Define preference as the fraction of signal attributable to host *i*:

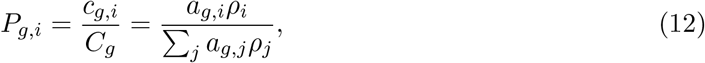

which is Eq. (3). This derivation remains valid under any monotone detection function provided preferences are defined by normalised weights of effective signal (the monotone transform cancels under normalisation).

## C Appendix C Multi-Phenotype Fitness Matrix and Three-Block Structure

### C.1 Wild fitness

Substitute Eq. (3) into Eq. (4):

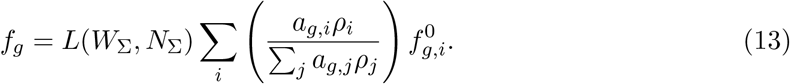

Let *C*_*g*_ = Σ _*j*_ *a*_*g,j*_*ρ*_*j*_. Then

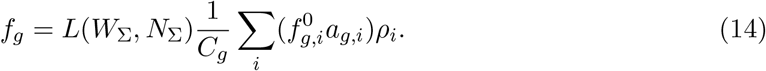

### C.2 Matrix form

Let ***f*** ∈ ℝ^*G*^ be the fitness vector over phenotypes, ***ρ*** ∈ ℝ^*m*^ host accessibility, and define matrices 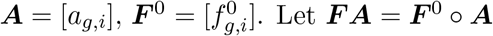 Define ***C*** = diag(*C*_1_, …, *C*_*G*_). Then

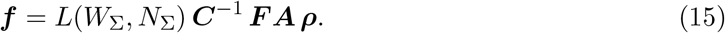

### C.3 Three-block interpretation

Equation (15) can be divided into three blocks as

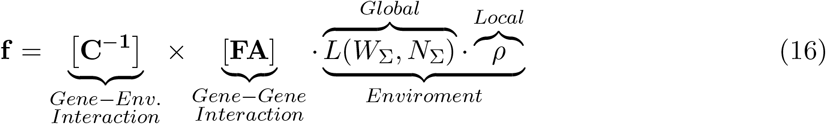

- **Environment block:** Describes global limitations *L*(*W*_Σ_, *N*_Σ_) and local landscape characteristics ***ρ***.
- **Gene–gene block: *F A*** = ***F*** ^0^ ◦ ***A*** Encodes the relationship between mosquito genes controlling olfactory sensitivity and feeding efficiency with host genes controlling emitted odour and nutritional value of blood.
- **Gene–environment block: *C***^*−*1^ Describes the relationship between genetically determined sensitivity and the attractant concentration determined by the local environment.

## D Appendix D Local Specialisation Index — Formal Construction and Bounds

### D.1 Cosine alignment

Define cosine similarity:

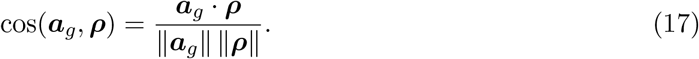

### D.2 Shannon effective diversity

For a nonnegative vector ***x*** = (*x* _1_, …, *x*_*m*_) with Σ_*i*_ *x*_*i*_ *>* 0, define proportions *p*_*i*_ = *x*_*i*_ Σ _*j*_ *x*_*j*_ and Shannon entropy *H*(***x***) = − Σ_*i*_ *p*_*i*_ ln *p*_*i*_. Define effective diversity (Hill number of order 1):

### D.3 Index and bounds

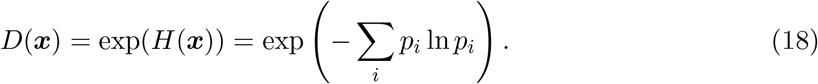

Define

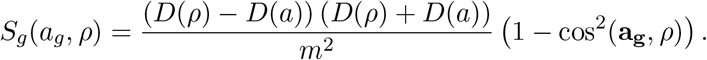

Since *cos*^2^(·) ∈ [0, 1] and *m* ≥ *D*(·) ≥ 1, the index is bounded. Under the scaling used above, one can show *S*_*g*_ ∈ [−1, 1] for typical ecological vectors; alternative normalisations can enforce the bound strictly (e.g. mapping the diversity ratio to [0, 1] via tanh). The interpretation in

Table 2 is preserved.

**E Appendix E: Coevolutionary Index — Rank-Based Interaction Complexity**

### E.1 Ranks

Let *r*_*a*_ = rank(***A***) and *r*_*f*_ = rank(***F*** ^0^). The maximal interaction complexity under independence is *r*_*f*_ *r*_*a*_. Let *r*_*F A*_ = rank(***F A***). If detection and exploitation are governed by shared latent factors (pleiotropy/constraint), ***F A*** becomes low rank relative to *r*_*f*_ *r*_*a*_.

### E.2 Index

Define

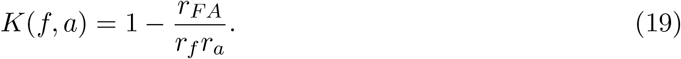

Then *K* ≈ 1 indicates strong constraint (low *r*_*F A*_), while *K* ≈ 0 indicates near-independence (*r*_*F A*_ ≈ *r*_*f*_ *r*_*a*_).

**F Appendix F: Fitness Expansion, Necessity of Correlation, and Simulation Recipe**

### F.1 Setup

For a single phenotype (drop subscript *g*), define random variables over hosts (with weights proportional to accessibility or preference as appropriate): *a, f* ^0^, and *ρ*. Let means 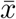, standard deviations *σ*_*x*_, and coefficients of variation 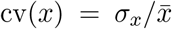. Define correlations corr(*x, y*) = Cov(*x, y*)*/*(*σ*_*x*_*σ*_*y*_) and co-skewness

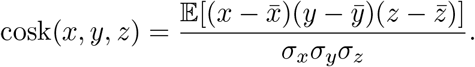

### F.2 Expansion (schematic form)

Starting from Eq. (4) with *P*_*i*_ ∝ *a*_*i*_*ρ*_*i*_, one can expand fitness around means to obtain

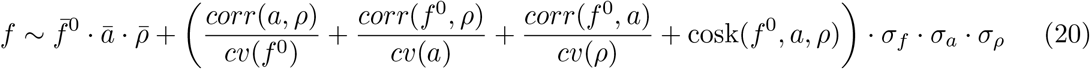

(Full algebra depends on the chosen weighting convention; the key point is that selective differentiation enters through correlation and higher-order terms.)

### F.3 Necessity of correlation for specialisation

If corr(*a, ρ*) = corr(*f* ^0^, *ρ*) = corr(*f* ^0^, *a*) = 0 and higher-order joint moments vanish, then Eq. (20) reduces to the baseline 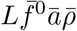. In that regime, selection cannot consistently favour host-selective strategies; fitness can only be improved by increasing means, consistent with generalism.

### F.4 Variability-regime logic (link to Figure 3 and Table 4)

When cv(*ρ*) is large, the term weighted by cv(*ρ*)^*−*1^ is suppressed, producing accessibility-driven selection. When cv(*ρ*) is small, intrinsic trait-variance terms dominate and the pathway depends on cv(*a*) vs. cv(*f* ^0^).

### F.5 Seasonality

Let *ρ*_*i*_(*t*) vary seasonally. If correlation signs in Eq. (20) reverse between seasons, selection can be dominated by the limiting season or by stable “refuge” hosts whose *ρ*_*i*_(*t*) is less variable.

### F.6 Simulation recipe for Figure 5

- A minimal simulation that reproduces the qualitative transition uses:
- Two hosts (*i* = 1, 2) with time-varying accessibility *ρ*_2_(*t*) increasing and *ρ*_1_(*t*) decreasing, e.g. relative accessibility *ρ*_2_*/ρ*_1_ *>* 1.
- Trait vectors *a*(*t*) = (*a*_1_(*t*), *a*_2_(*t*)) and 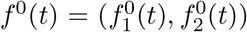 updated by a simple selection gradient:

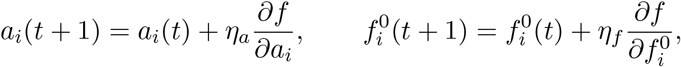

with small step sizes *η*_*a*_, *η*_*f*_ and bounds enforcing positivity.
- Genetic coupling implemented by projecting updates onto a low-dimensional manifold (high-*K* case) versus allowing independent updates (low-*K* case).
- At each *t*, compute *P*_*i*_(*t*) from Eq. (3), compute *S*_*g*_(*t*) from Eq. (6), and compute *K*(*t*) from Eq. (7) (or from the evolving matrices in a multi-phenotype version).

Plot *ρ*_*i*_(*t*), *P*_*i*_(*t*), *S*_*g*_(*t*), and *K*(*t*) to obtain Figure 5.

#### Practical note

In practice, Figure 5 can be generated either from an explicit multi-phenotype matrix simulation (preferred for computing rank-based *K* directly) or from a stylised coupling proxy that tracks an “effective” *K*(*t*).

